# On the onset of multicellular invasive behavior in hierarchical lineage: the role of inhibitory feedback and local fluctuations

**DOI:** 10.1101/2025.06.17.660235

**Authors:** Josué Manik Nava-Sedeño, Abraham Martínez, Haralampos Hatzikirou

## Abstract

The emergence of multicellular invasive behavior is a key characteristic of various biological processes, including wound healing, development, and tissue regeneration. In this study, we develop a lattice-gas cellular automaton (LGCA) model to explore the role of inhibitory feedback in the invasive behavior of a hierarchical lineage composed of stem cells and differentiated cells. We consider both non-spatial and spatial stochastic models to investigate how spatial interactions influence invasion dynamics. Our findings suggest that inhibitory feedback from differentiated cells significantly impacts the invasive potential of stem cells. In addition, local fluctuations induce unstable fronts that move with relatively low speed. Finally, we explore the implications of our work for understanding the regulation of multicellular dynamics in various pathophysiological contexts.

## 1. Introduction

Multicellular invasion is a fundamental process that underlies various biological phenomena, including wound healing, tissue development, and regeneration (1; 2; 3; 4; 5). The hierarchical organization of cell populations, often comprising a small subpopulation of stem cells and a majority of differentiated cells, has been shown to play a critical role in the regulation of growth and invasiveness (6; 7; 8; 9; 10). Stem cells possess the unique ability to self-renew and differentiate, thus driving both tissue expansion and the maintenance of cellular heterogeneity. Conversely, differentiated cells are nonproliferative but can exert inhibitory effects on stem cells, potentially restricting their self-renewal and limiting overall growth (11; 12). The dynamics of these interactions are highly dependent on the spatial organization of cells, which can significantly influence the emergence of invasive behavior (13; 14).

In the context of pathology, tumors represent a prime example of aberrant multicellular invasion. Tumors can be classified as malignant or benign, depending on whether they invade surrounding tissues or remain confined to their tumor mass. The degree of malignancy often correlates with the speed of invasion into neighboring tissues; faster invasion is typically associated with higher malignancy. This process is often marked by the appearance of “invasive fingers”, irregular protrusions along the tumor boundary where some regions advance more rapidly than others, forming structures resembling fingers (15; 16). Tumor malignancy and the formation of fingering patterns during cell invasion have been heavily studied through biophysical (17; 18) and mathematical modeling (19; 20; 21; 2), especially employing partial differential equations for finding the growth speed of tumor spheroids (22; 23), and instability conditions of spatially homogeneous solutions (24).

In contrast, the hierarchy of tumor lineages, as well as inhibitory interactions, has been widely considered in mathematical models to explain the onset of heterogeneity, tumor recurrence, selection of therapy-resistant phenotypes, and tumor response to therapy, among others (25; 26; 27). Furthermore, the effect of feedback on cell lineages leads to spatial patterning, in particular in physiological conditions (28). However, tumor cell hierarchy and inhibitory interactions between them have not yet been investigated as key mechanisms of malignancy and tumor roughness formation.

To better understand the interplay between stem cells and differentiated cells in the context of multicellular invasion, we propose a lattice-gas cellular automaton (LGCA) model that accounts for both reactive cellular kinetics and spatial interactions. LGCA models have been successfully used to describe various aspects of multicellular dynamics, providing a discrete framework that allows for the modeling of individual cell behavior, including proliferation, differentiation, apoptosis, and migration (29; 30; 31; 32; 33). Here, we extend the LGCA approach to study the effects of inhibitory feed-back from differentiated cells on stem cell self-renewal and differentiation, and how these effects shape the invasive potential of multicellular systems.

We present a series of models, ranging from a non-spatial Markov chain model to spatial LGCA models in one and two dimensions. By analyzing the stability of these models and performing extensive numerical simulations, we aim to characterize the conditions under which invasive behavior emerges as a function of inhibitory feedback and local fluctuation. Our results provide new insights into the role of hierarchical lineage interactions in multicellular invasion and suggest potential avenues for therapeutic intervention targeting stem cell differentiation and inhibition.

## 2. Model definition

We consider a tumor cell population with a hierarchical differentiation lineage. To simplify this hierarchy and keep the model mathematically tractable, we reduce the hierarchical lineage to only two cell populations: cancer stem cells, which can divide indefinitely and are immortal, and totally differentiated cells, which cannot reproduce and are not immortal but can exert pressure on cancer stem cells, affecting their self-renewal and differentiation behavior. Please note that with the term stem cells, we denote all proliferating lineage populations, which include stem and progenitor cells. To study the effect of spatial dimensionality on tumor dynamics, we consider three different models: a non-spatial model, or 0D model, consisting of a discrete Markov chain, and two spatial models, 1D and 2D models, formulated as LGCA.

### 2.1. Temporal Markov chain model

For the non-spatial model, we consider that all cells can interact with each other, i.e., the interaction network is a complete graph. Both stem (*S*) and terminally differentiated (*T*) cells undergo self-renewal, differentiation, apoptosis, or quiescence in discrete time steps. The interactions between both cell populations during these processes and their mathematical abstraction in the model are depicted in Fig. 1 Furthermore, we assume that at every time step, only one of these four possible processes occurs. We also consider that the total number of stem cells *n*_*S*_ and terminally differentiated cells *n*_*T*_ cannot increase above a maximum of *K*_*S*_ and *K*_*T*_ stem and terminally differentiated cells, respectively, due to limited space and resources in the tumoral environment.

**Figure 1.**
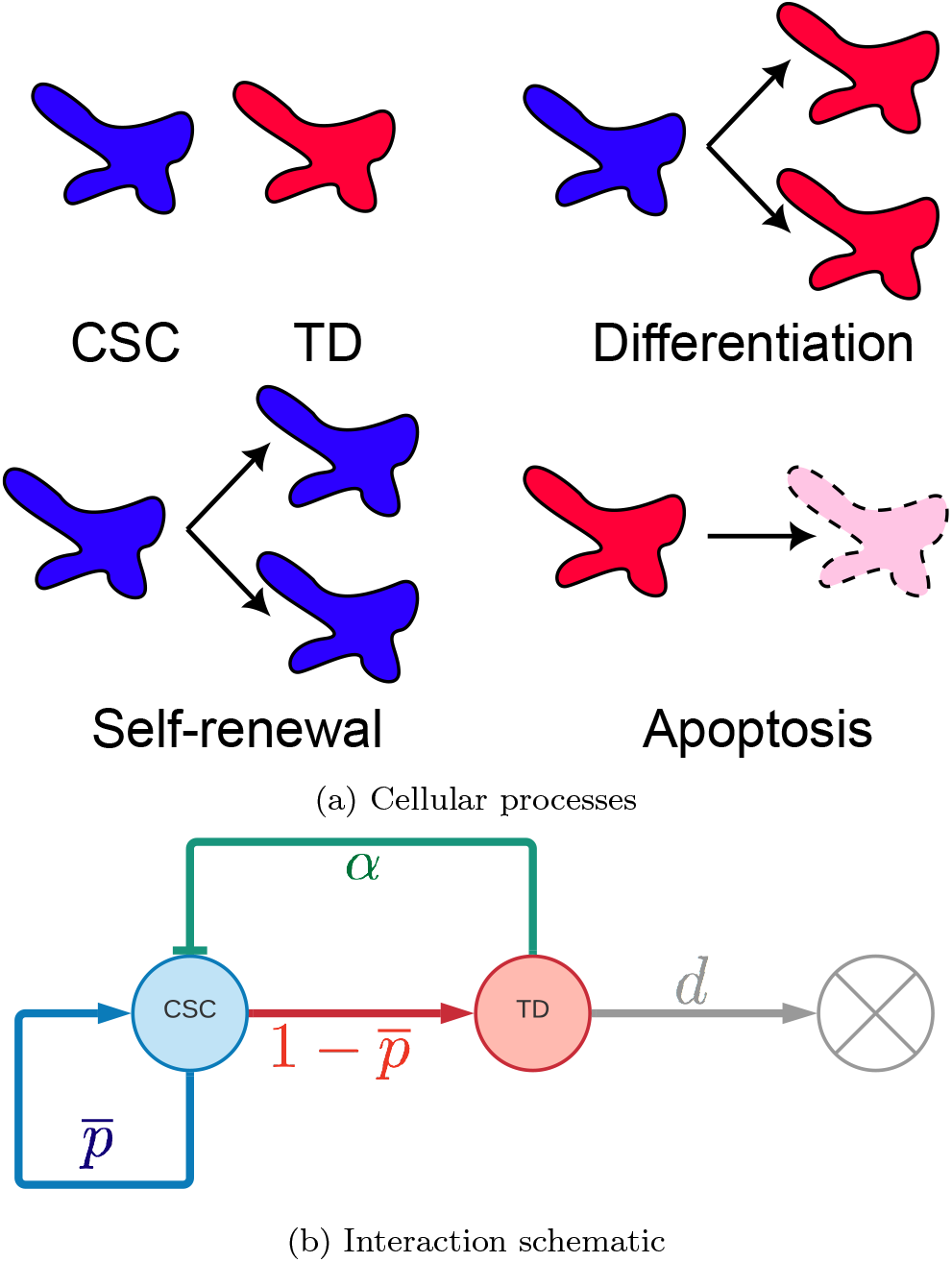
Modeling of cancer cell dynamics. 1a. Cellular processes considered in the model. We assume two cell phenotypes: cancer stem cells (CSC) and totally differentiated cancer cells (TD). CSCs can differentiate to produce two TDs, CSCs can self-renew to produce two CSCs, and TDs can undergo apoptosis. Additionally, cells remain quiescent in the absence of these processes. 1b. Schematic of the mathematical modeling of cell processes in our models. Terminally differentiated cells can either stay quiescent, or undergo apoptosis at a rate *d*. In the absence of terminally differentiated cells, stem cells self-renew with probability 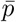, and differentiate with probability 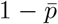 whenever they divide. Terminally differentiated cells, however, have an inhibitory feedback effect on stem cells, affecting their probability to self-renew or differentiate by an exponent *α*.

#### Self-renewal

During this process, a stem cell divides symmetrically, thus increasing the number of stem cells by one. We assume that only one stem cell is able to self-renew, as long as at least one stem cell is present and as long as the number of stem cells is maintained at or below *K*_*S*_. We assume that self-renewal happens at a rate *ν*, that the selected stem cell can choose between self-renewal and differentiation with probability *p* and 1 − *p*, respectively,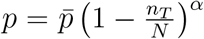, where 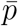 is the probability of self-renewal in the absence of terminally-differentiated cells, *N* := min {*K*_*S*_, *K*_*T*_} is a parameter indicating the maximum sustainable number of cells of each phenotype per lattice site, and *α* is a parameter denoting the strength of the feedback inhibition of terminally-differentiated cells on stem cells. This inhibitory feedback broadly models contact-mediated and paracrine-based negative control of stem cell self-renewal. The logistic term 1 − *n*_*T*_ */N* can be viewed as contact inhibition when *α* = 1. However, for *α >* 1 this inhibition is enhanced by additional biological mechanisms such as paracrine signals (e.g., TGF*β*) that inhibit stem cell renewal. The probability of a self-renewal event happening is thus given by

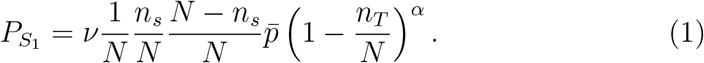

In general terms, this expression considers choosing CSC randomly and uniformly, then choosing an empty CSC site for the new daughter cell to be placed in, as well as the probability that the selected stem cell underwent self-renewal.

#### Differentiation

During this process, a stem cell divides producing two terminally differentiated cells, thus decreasing the number of stem cells by one, and increasing the number of terminally differentiated cells by two. As with self-renewal, we assume a stem cell is picked randomly for differentiation given there is at least one stem cell present, and as long as two terminally-differentiated cells are sustainable. Differentiation happens at a rate *ν* as well, with probability 1 − *p*. Therefore, the probability of a differentiation event happening is given by

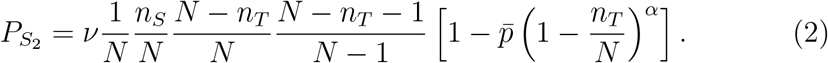

In this case, this is the probability of choosing a CSC randomly and uniformly, then choosing two empty TD sites for the new differentiated daughter cells to be placed in, and considering the probability that the selected stem cell underwent differentiation.

#### Apoptosis

Terminally differentiated cells cannot proliferate nor differentiate, but can undergo apoptosis at a rate *d*. During this process, the number of terminally differentiated cells decreases by one. We assume apoptosis happens to a randomly selected terminally-differentiated cell, as long as there is at least one terminally-differentiated cell. Thus, the probability of an apoptosis event happening is given by

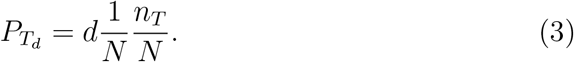

This equation considers the probability that a TD cell is randomly and uniformly chosen, and that independently, the chosen cell undergoes apoptosis.

#### Quiescence

Finally, all cells of both phenotypes may remain quiescent. This happens if no other process happens, and the number of cells of both types remains unchanged, i.e.

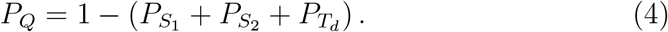

This expression is just the probability of the complement process to self-renewal, differentiation, and apoptosis; namely, the only other process which can happen every time step: quiescence.

### 2.2. Spatial LGCA models

For spatially structured tumors, we define an LGCA extension of the discrete Markov chain process explained previously. In the LGCA framework, single cells are abstracted as point particles interacting and moving on a discrete lattice. As such, cells are characterized by both their spatial position and velocity. More formally, space is divided into a discrete lattice, ℒ *⊂* ℝ. Here, we consider both a one-dimensional lattice ℒ= ℤ, and a two-dimensional square lattice ℒ= ℤ^2^. Cell positions are described by their location 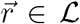. Cells on the lattice may only move to neighboring lattice sites. As such, in 1D cells may have *b* = 2 different non-zero velocities, *c*_1_ = 1 and *c*_2_ = − 1, while in a 2D square lattice, cells may have *b* = 4 different non-zero velocities, 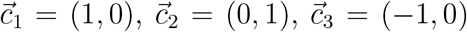, and 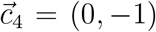. Additionally, cells may rest and remain stationary. These cells have a zero velocity, *c*_0_ = 0 and 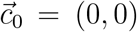 in 1D and 2D, respectively. We indicate the number of cells with velocity 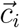 (or *c*_*i*_ in 1D) through the occupation number *s*_*i*_. Furthermore, we impose an exclusion principle on the cells, such that no more than one cell may move in the same direction within the same lattice site, and no more than *a ∈* ℕ_0_ cells may rest in every lattice site, where *a* is a free model parameter. Thus, we consider *s*_0_ *∈* {0, …, *a*}, and *s*_*i*_ *∈* {0, 1}, for *i* ≠ 0. Consequently, the number of cells in a lattice site 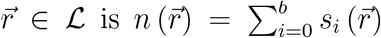, up to a maximum of *K* = *a* + *b* cells per lattice site. In our model, we consider two separate lattices, one for the stem cell population, and a second one for the terminally differentiated cell population. Each lattice will consider a different number *a*_*γ*_, *γ ∈ S, T* of resting cells, resulting in a different maximum number of cells per node for each lattice, *K*_*S*_ and *K*_*T*_ . We refer to the vector of occupation numbers and the cell number per lattice site of stem cells and terminally differentiated cells by 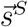 and 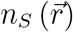, and 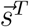 and 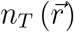, respectively. Moreover, we denote the total number of cells for both populations per site as *n* = *n*_*T*_ + *n*_*S*_. Note that the exclusion principle only applies to cells of the same type, i.e. it is possible for a stem cell and a differentiated cell to have the same position and velocity.

At every time step, three operators, reaction ℛ, reorientation 𝒪 and translocation 𝒯, are applied sequentially to every node in the lattice synchronously, after which we consider that a single time step has elapsed.

At the beginning of the time step, the reaction operator ℛ is applied to every node independently and synchronously. This operator applies one step of the non-spatial Markov model to the node, i.e. it increases or decreases the number of stem and/or terminally differentiated cells by calculating the probabilities 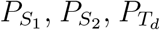 and *P*_*Q*_ for the number of *S* and *T* cells in the node, determining which of the four processes will occur according to these probabilities, picking one cell within a velocity channel randomly and uniformly to undergo this process and, in the case of self-renewal and differentiation, picking empty channels randomly and uniformly in the appropriate lattice to be occupied by the daughter cells.

After any of the four reactive processes have occurred in every lattice site, the reorientation operator 𝒪 is applied independently and synchronously to all nodes. The operator redistributes cells randomly and uniformly among all velocity channels to simulate random changes in velocity during the diffusive motion of cells with no external directional cues. This is done through the uniform transition probability.

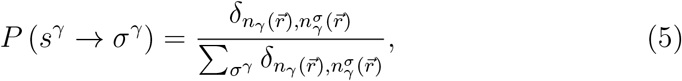

where *γ ∈* {*S, T*} refers to the stem cell or terminally differentiated lattices, *σ*^*γ*^ is the vector of occupation numbers of one of the *γ* lattice after velocity reorientation, *δ*_*i*,*j*_ is the Kronecker delta, which ensures that cell number is conserved after cells change movement direction, and 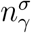 is the number of cells after cells reorient. After both reaction and reorientation operators have been applied, we denote the occupation number of the *i*-th velocity of the *γ* lattice site *r* by 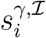, the post-interaction occupation number.

Finally, to simulate cell movement, the translocation operator is 𝒯 applied, which deterministically moves cells from their present node to the neighboring node in the direction of their velocity. If we assume that, before reaction, reorientation, and movement, the time step is *k*, then we say that after all these processes have taken place, the time step has increased by one to *k* + 1. Therefore, the dynamics can be summarized by the stochastic difference equation

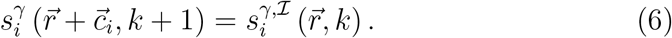

A summary of the model parameters, their biological meaning, and typical values used in simulations are presented in Table 1.

**Table 1.** Parameters of the model, their biological interpretation, and typical values used for computer simulations.

| Parameter | Meaning | Values |
| --- | --- | --- |
| $K_S$ | CSCs size/compressibility | 8 |
| $K_T$ | TDs size/compressibility | 16 |
| $\nu$ | CSC mitosis rate | 0.8 |
| $\bar{p}$ | Ratio of self-renewal events to total proliferative events in absence of TDs | 0.7 |
| $\alpha$ | TD inhibitory pressure | 2 |
| $d$ | Apoptosis rate | 0.5 |
| $1/\mu$ | Average duration of the cell cycle | 0.1 |

### 2.3. Cell motility

The dynamics of the nonspatial (0D) and LGCA (1D and 2D) models is illustrated graphically in Fig. 2. In order to characterize cell motility with respect to the spatial models’ parameters, we recall that at every node, up to *b* cells may move to a neighboring node, while *a* cells may remain stationary. Furthermore, we rescale the automaton such that every lattice site is located a distance *E >* 0 with respect to its first nearest neighbors, and such that time increases by *τ >* 0 every time step. Since cells reorient uniformly and independently, the probability of finding a certain cell is given by

**Figure 2.**
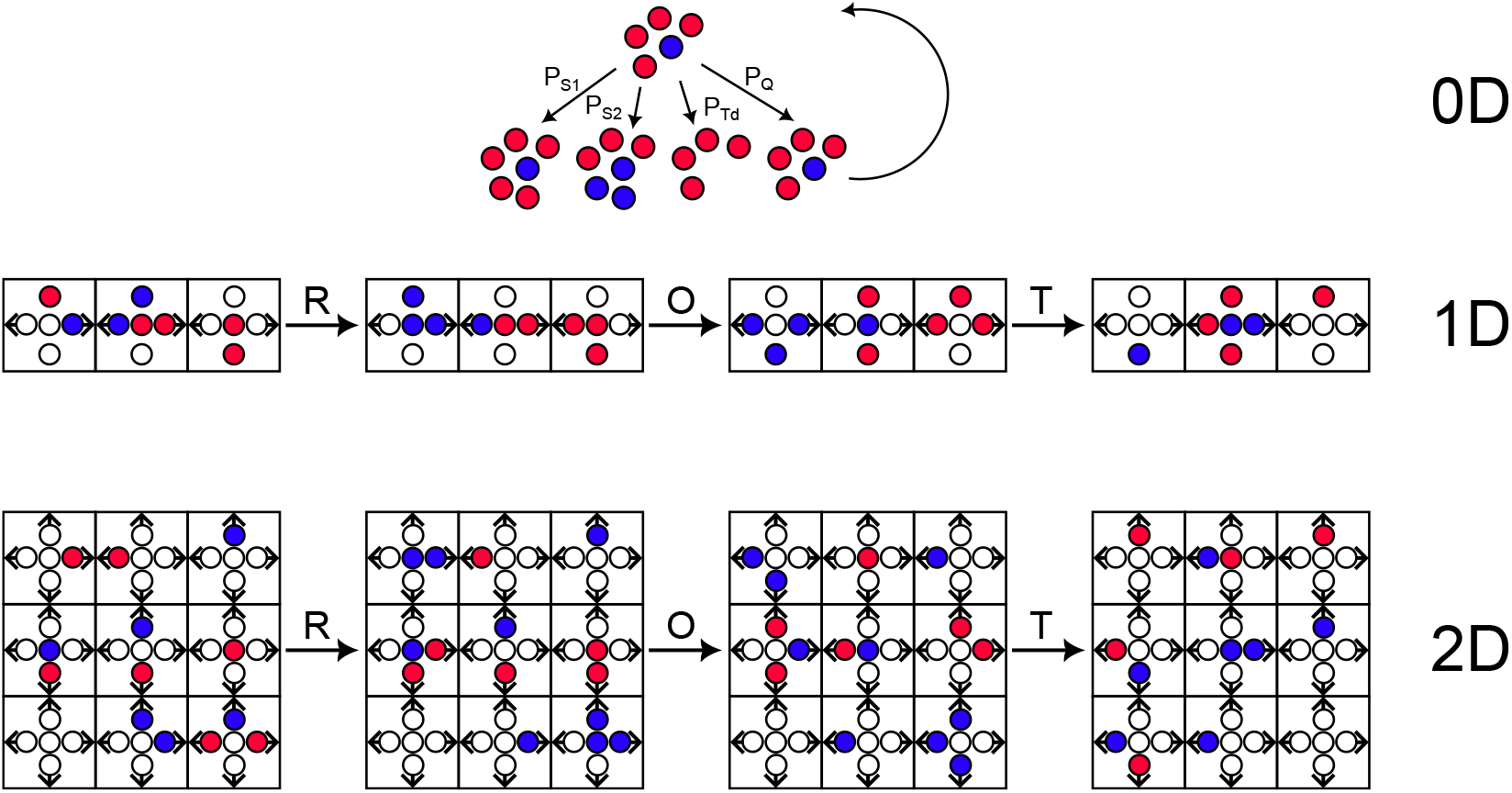
Schematic of the model dynamics. In the non-spatial stochastic process model (0D), cells at most one cell of the entire population can undergo self-renewal, differentiation, apoptosis, or quiescence with probabilities 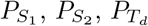, or *P*_*Q*_, respectively. After one of these processes has occurred, it is considered that one time step has elapsed, and the process repeats for as many time steps as desired. In the one-dimensional LGCA model (1D), every lattice site, in parallel, undergoes one of the aforementioned processes with their respective probabilities, as long as there are velocity channels available. This is the reaction (ℛ) step. Afterwards, all cells in every lattice site are randomly and uniformly assigned a new velocity channel, to simulate cellular diffusion. This is the reorientation (𝒪) step. Finally, all cells are moved to the left or right neighboring sites, or they remain on their own site, depending on their velocity channel, to simulate cell migration. This is the translocation (𝒯) step. After all four steps have occurred, a time step has elapsed and the process begins again. In the two-dimensional LGCA model (2D), the process is identical, except that cells can now also move upwards and downwards.

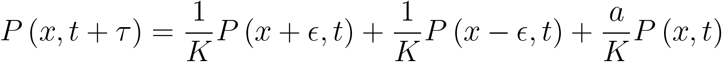

and

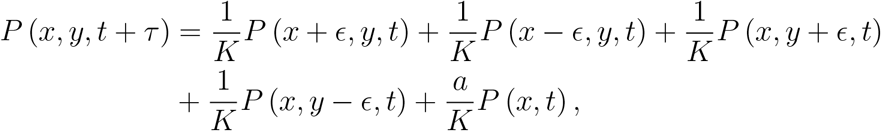

for the one-dimensional and two-dimensional case, respectively, where *x, y*, and *t* are the rescaled, continuous spatial and temporal variables. Letting *E →* 0 and *τ →* 0 under diffusive scaling, we find that 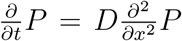 and 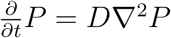, for the 1D and 2D case, respectively, where in both cases, the diffusion coefficient is given by

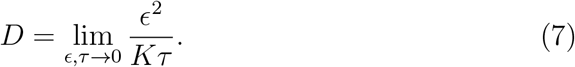

Thus, we see that the cell motility is inversely proportional to the maximum number of cells per node *K*. Since *K* = *a* + *b*, and *b* is fixed by the lattice geometry, we can control the motility of each cell phenotype through *a*, the maximum number of resting cells per node, since stem and terminally differentiated cancer cells have different migratory behavior (4).

## 3. Results

### 3.1. Cell population kinetics: Non-spatial model

We first consider the on-node reactive dynamics of the model without spatial considerations, i.e. without considering the effects of cell migration. Given that the effect of the reaction step is only increasing or decreasing the total number of cells, the stochastic difference equations describing the dynamics of the non-spatial model are given by

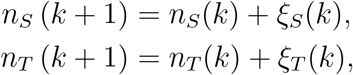

where *ξ*_*S*_(*k*) and *ξ*_*T*_ (*k*) are independent and identically distributed random variables such that *ξ*_*S*_(*k*) *∈* {−1, 0, 1} and *ξ*_*T*_ (*k*) *∈* {−1, 0, 2} and

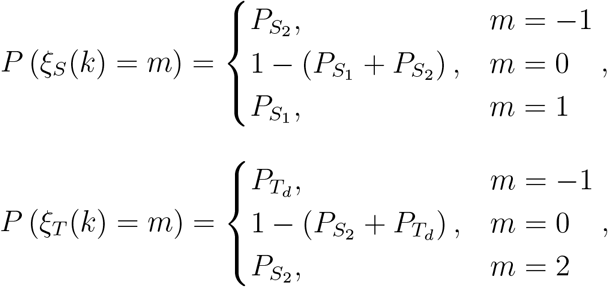

for all *k ∈* ℕ. Taking the expected value on both sides of the stochastic finite difference equations, and under a mean-field approximation such that *(f* (*X, Y*)) = *f* (*(X)*, *(Y)*), we obtain the system of finite difference equations

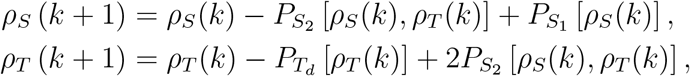

where *ρ*_*γ*_(*k*) := *n*_*γ*_(*k*) . Rescaling the automaton with a small time step length *τ*, and subtracting *ρ*_*S*_(*t*) and *ρ*_*T*_ (*k*) from the first and second equation, respectively, we obtain the approximate ordinary differential equation (ODE) system

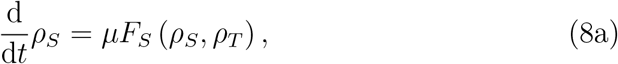

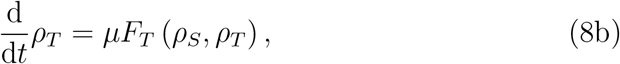

where 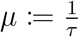 is the reaction rate dependent on the automaton scaling, and

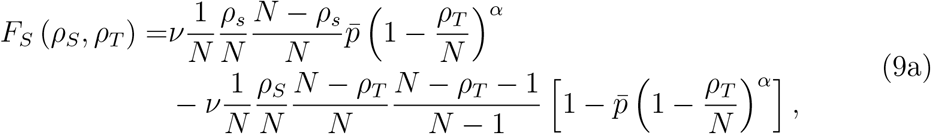

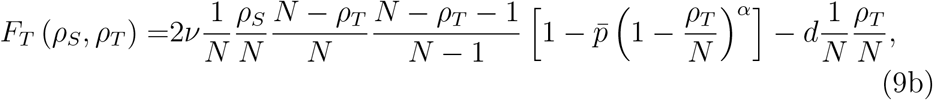

are the reactive terms. Note that *ν* can be factored out of Eqs. 9. Thus, rather than *ν* itself, important parameters in the model are the product *µν* and the ratio 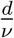.

#### 3.1.1. Linear stability analysis

The mean field ODE approximation, Eqs. 9, has two biologically relevant steady states, the trivial steady state *ρ*_*S*_ = *ρ*_*T*_ = 0, corresponding to the extinction of both populations, and a non-zero steady-state 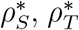 which has no closed form, representing the long-term survival and coexistence of both phenotypes.

Linearizing about the trivial steady state yields the Jacobian matrix

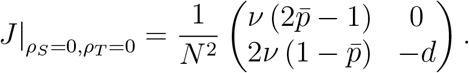

Recalling the stability criteria for two-dimensional systems, det(*J*) *>* 0 and tr(*J*) *<* 0, and looking at the determinant of the Jacobian matrix, we find that the trivial steady state is an unstable saddle point when

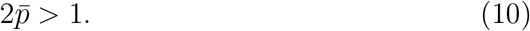

To determine the stability of the steady state outside of the regime, we turn to the trace of the Jacobian matrix, which gives the condition

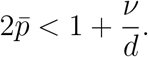

Since *ν* and *d* are both strictly positive parameters, we see that whenever 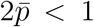 (the regime where the trivial steady state is not a saddle point), necessarily 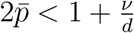, since 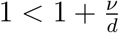. Thus, the steady state with no cells of either phenotype is stable when 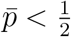. Biologically, this condition tells us that, in the absence of space, cells die out only if stem cells differentiate more often than self-renew when in isolation. A numerical linear stability analysis of the non-trivial steady state reveals it gains stability in the range given by Eq. 10 through a transcritical bifurcation, as evidenced by the bifurcation diagram (Fig. 3).

**Figure 3.**
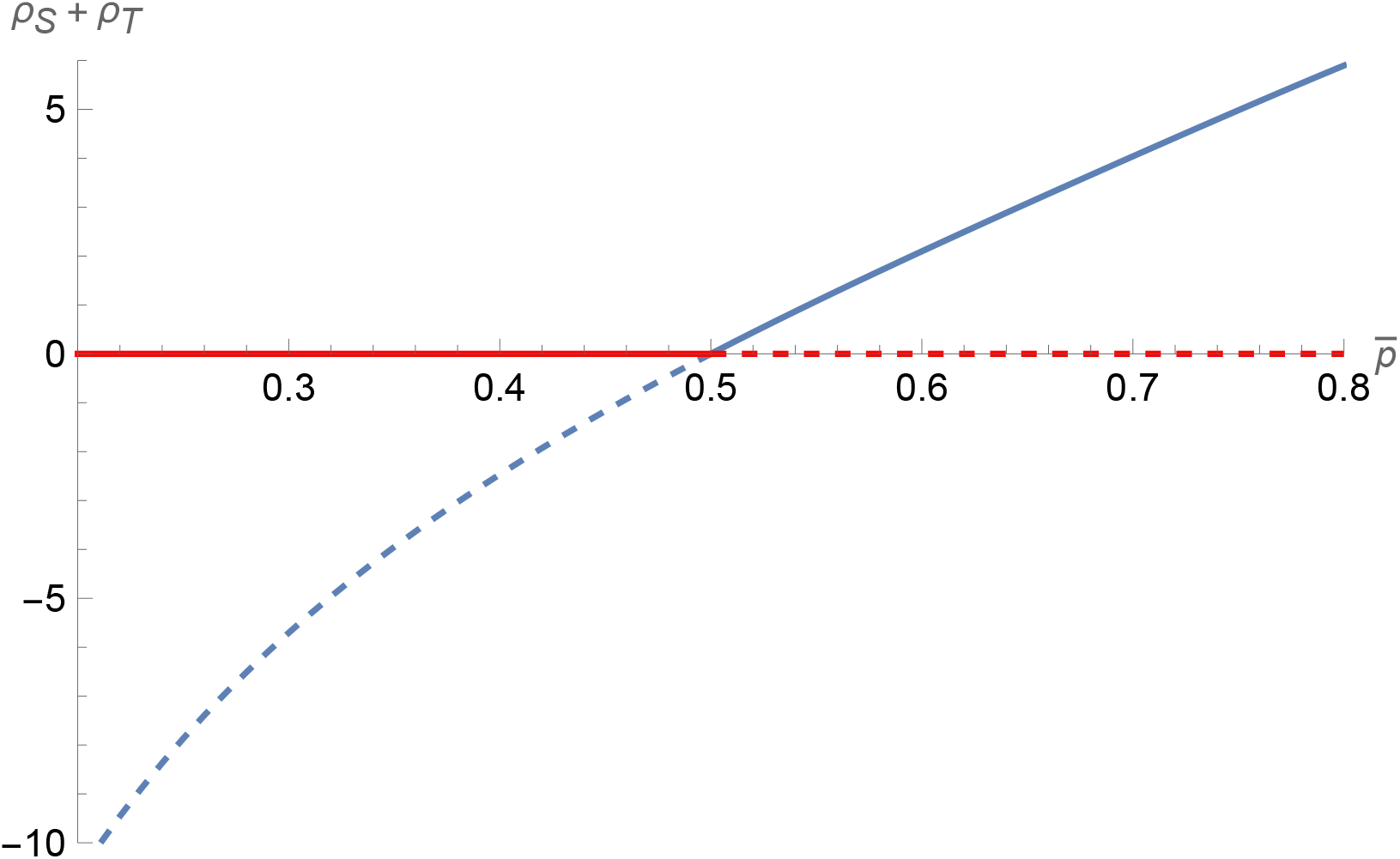
Bifurcation diagram of the approximate non-spatial ODE model. The system experiences a transcritical bifurcation at 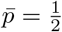. Red lines show the extinction state, blue lines the coexistence state. Solid and dashed lines indicate stable and unstable states, respectively. Parameter values were set to *ν* = 0.8, *d* = 0.1, *N* = 10, and *α* = 2.

It should be noted, however, that the cell-free state is an absorbing state of the original Markov chain, and therefore the process will always end up at this trivial state almost surely, while the mean-field ODE system will asymptotically approach the non-trivial steady state whenever Eq. 10 is not fulfilled. Nonetheless, the mean-field ODE approximation describes the expected behavior of the Markov chain model quite accurately for short and medium time spans. As show in Fig. 4, in the 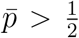 regime, it may take a very long time to observe significant differences between the mean of the process and the ODE approximation, in this case, more than 1 *×*10^3^ time steps for a Markov chain with only nine states (corresponding to a maximum number of eight cells). The mean time to extinction grows quickly with the maximum number of cells.

**Figure 4.**
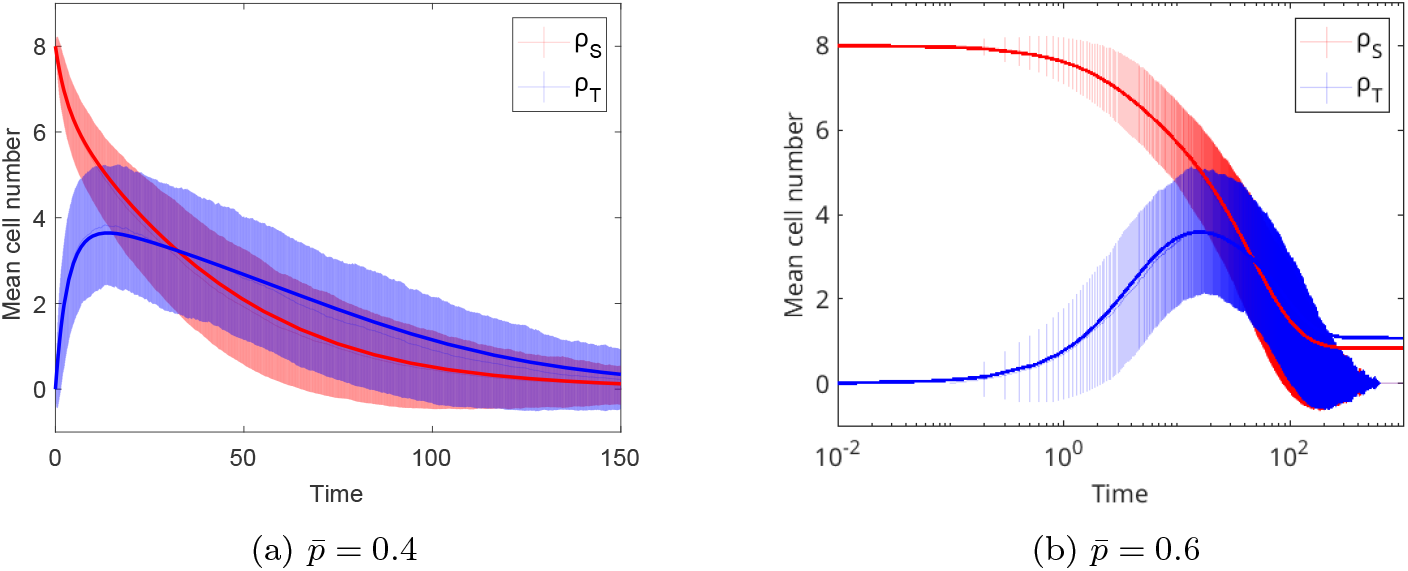
Non-spatial stochastic model and ODE approximation. Solid lines show numerical solutions to Eqs. 8, shaded areas correspond to one standard deviation around the sample mean of the stochastic process obtained from 2 *×* 10^3^ realizations. The value of 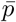 was set at 0.4 (left) and 0.6 (right). Other parameter values were set at *ν* = 0.8, *α* = 2, *d* = 0.5, *N* = 8 and *τ* = 1*/µ* = 1*/*10. As observed, the ODE approximates the mean of the stochastic process accurately at all times for 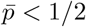, while the approximation completely separates from the mean of the process for 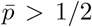 in the long-time limit. Error bars correspond to one standard deviation.

Additionally, we numerically determined the fraction of stem cells (with respect to the total cell population) at the steady state of Eqs. 8 for changing values of the self-renewal 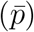, differentiated cell death (*d*), and inhibition feedback (*α*) parameters. As seen in Fig. 5, the higher the inhibition feedback strength, the smaller the region in parameter space where stem cells are plentiful, which highlights the importance of this negative feedback loop for keeping the population in check. Naturally, lower self-renewal probabilities also result in lower stem cell fractions. However, at intermediate values of self-renewal probability and inhibition feedback strength, the effect of cell death of differentiated cells is non-monotonic on the fraction of stem cells, as higher fractions of stem cells result from both low and high number of cell death parameter values. This may be due to the fact that, at small death rates, stem cells more often undergo self-renewal, since all cellular processes are mutually exclusive, while at high death rates, stem cells are less inhibited to undergo self-renewal.

**Figure 5.**
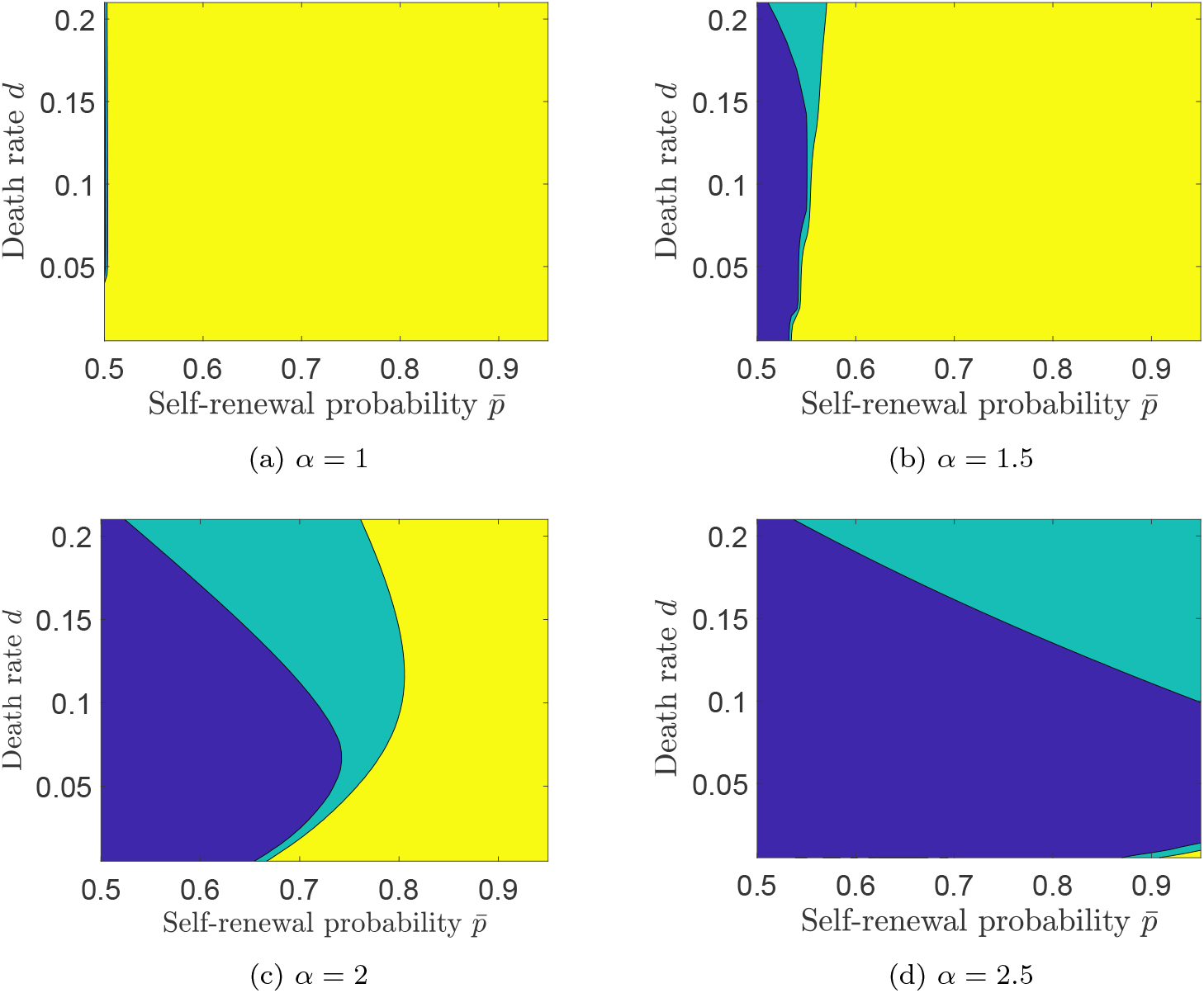
Stem cell fraction over the whole cell population at the steady state. Blue corresponds to regions in parameter space where the steady state stem cell fraction is below 10% of the whole cell population, turquoise indicates stem cell fractions between 10% and 20%, and yellow corresponds to stem cell fractions over 20%. Other parameter values were fixed at *ν* = 2 and *N* = 8.

### 3.2. Invasion dynamics in one dimension

We now consider the full dynamics of the model in a 1D setting corresponding to the invasion of cells in a restrictive environment, such as that of tissues with a high ECM density, or in a highly structured tissue, such as colon crypts, assuming that a solid tumor is located at the left, such that cell movement is restricted in the left direction. We considered a semi-infinite lattice, ℒ = ℕ, with reflecting boundary conditions at the boundary, i.e., cells that would normally cross the boundary will instead have their velocity inverted to prevent their escape. We considered an initial condition where the node at the boundary has *N* = min {*K*_*S*_, *K*_*T*_} cells, and all other nodes are empty, in both stem and terminally differentiated cell lattices.

When considering the complete model dynamics, we observe that the reactive dynamics considered previously still take place at every point in space simultaneously. Additionally, after the reaction occurs, cells now move randomly to a neighboring position or rest. If we now define the post-reaction cell density as

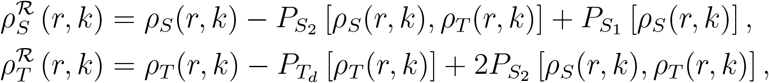

which are just the cell densities after one time step in the non-spatial approximation, we see that in the full one-dimensional approximation, the discrete dynamics of the model are given by

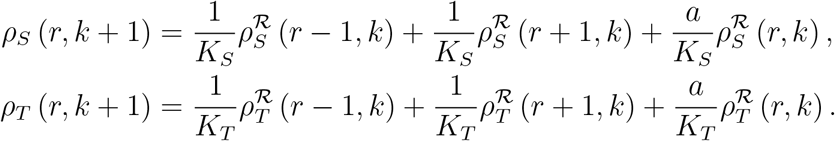

Since only one reaction event can take place every time step, we can assume that spatial fluctuations of the post-reaction cell density are mainly due to spatial fluctuations of the pre-reaction cell density, i.e., we may approximate

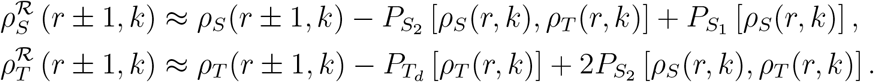

Substituting these approximations into the difference equations, we obtain

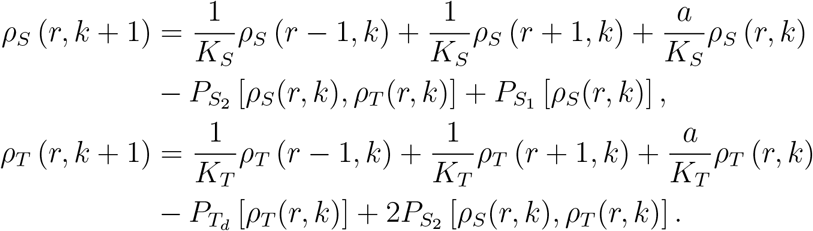

Rescaling the system of difference equations with a small time step *τ* and lattice size *E* and subtracting *ρ*_*S*_ (*x, t*) and *ρ*_*T*_ (*x, t*) from both sides of the equation, we arrive at the reaction-diffusion PDE system

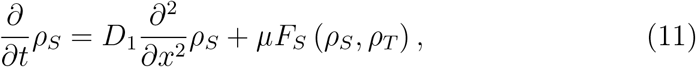

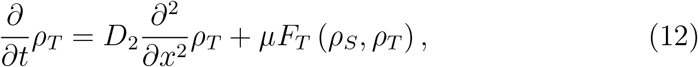

where the reaction rate *µ* and the reaction rates *F*_*S*_ and *F*_*T*_ are defined as before, and the diffusion coefficients are given by 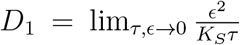 and 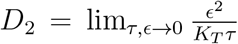, as found before, where *K*_*S*_ and *K*_*T*_ are the maximum number of *S* cells and *T* cells, respectively. Thus, by choosing different values for the maximum number of resting cells *a*_*γ*_ for each phenotype, we are able to capture differences in motility between phenotypes.

#### 3.2.1. Speed of the invasive front

To investigate the effect of cellular processes and interaction on the invasive speed of the tumor, we assume that, at *t* = 0, the tumor is confined to the main tumor body, which we set at *x* = 0, and for *t >* 0 cancer cells start invading the surrounding healthy tissue to the right of the tumor (*x >* 0). Thus, we assume that non-homogeneous solutions are traveling waves, i.e., they depend on the independent variables solely through the relation *z* = *v* − *vt*, where *v* is the speed of the invasive front to be determined. Using a change of variables, we can transform the PDE system into an ODE system, noting that 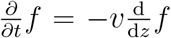 and 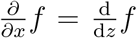 using the chain rule, where *f* (*x, t*) = *f* (*z*) is the traveling wave solution to Eqs. 11 and 12. Using this change of variables, we can rewrite Eqs. 11 and 12 as a system of two second-order ODEs

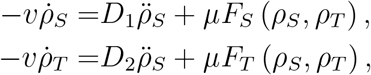

Where 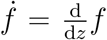. Defining 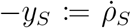 and 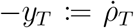 we obtain a system of four first-order ODEs

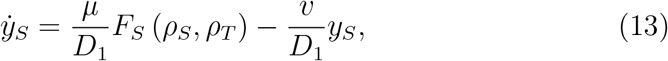

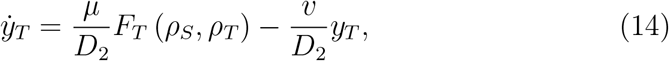

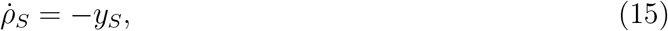

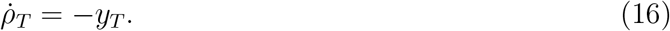

From Eqs. 9a and 9b, we note that *ρ*_*S*_(*z*) = 0, *ρ*_*T*_ (*z*) = 0, *y*_*S*_(*z*) = 0, *y*_*Z*_(*z*) = 0 is a steady-state solution of the ODE system. Linearizing the system around this steady state yields the Jacobian matrix

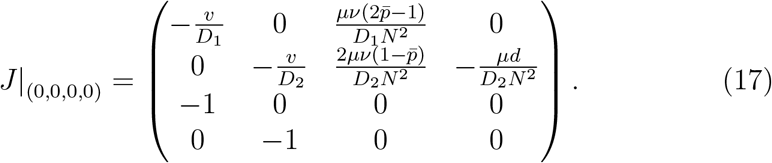

The Jacobian matrix has two real eigenvalues with opposite signs, indicating saddle point behavior, and two eigenvalues that could be real or complex conjugates. If these eigenvalues were complex conjugates, the steady state would be a spiral. However, oscillations near the zero steady state would result in a traveling wave with negative *ρ*_*S*_ and *ρ*_*T*_ values, which are not biologically realistic. Therefore, we impose the condition that all eigenvalues are real, which yields the condition for the minimum wave front speed.

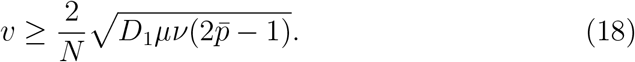

Two important features are noteworthy. First, when 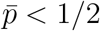, the inequality is not well defined; thus, one may argue that traveling wave solutions only exist in the parameter range where the non-zero steady state is stable according to the non-spatial ODE model. Second, the minimum front speed does not depend on *D*_2_, the terminally differentiated cell diffusion coefficient; thus, the speed of the invasive front only depends on the motility of stem cells, not differentiated cells. Additionally, greater reaction *µ* and division *ν* rates also increase the speed of the invasive front.

Since the inhibition feedback does not contribute linearly to the front propagation speed, we investigated its nonlinear effects by simulating theone-dimensional LGCA model. We considered the case 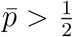, where the tumor population propagates instead of quickly dying out. Computationally, we calculated the ensemble average of the stem cell and terminally differentiated cell populations as a function of space, at two different timepoints and for two different values of the inhibition feedback parameter *α*. As observed in Fig. 6, the invasive front speed (observed as the displacement of the front between two fixed time points) does not seem to be significantly affected by the different values of *α*, in agreement with the linear analysis performed previously. However, at low values of this parameter, stem cells greatly outnumber terminally differentiated cells, and the total number of tumor cells is considerable. On the other hand, when *α* is high, both stem cell and terminally differentiated cell populations are commensurate, and the population size per site is, on average, of just a couple of cells, which makes the tumor much more susceptible to extinction due to random fluctuations.

**Figure 6.**
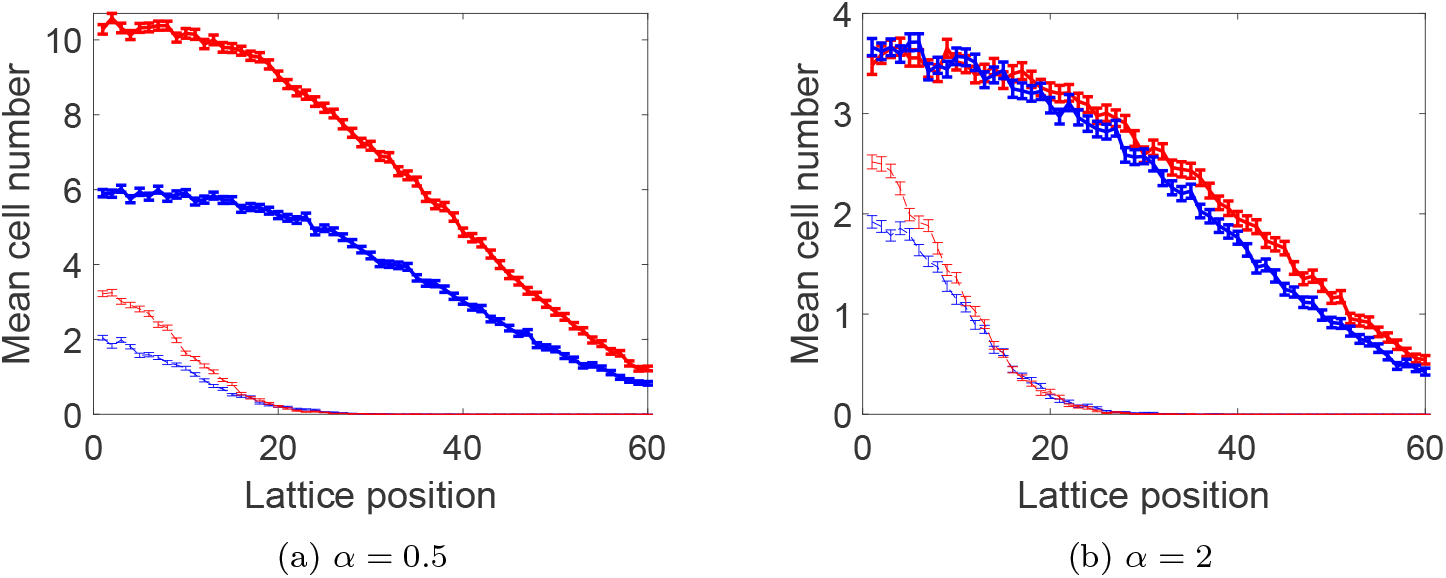
One-dimensional LGCA simulations. Blue and red lines indicate the ensemble average of terminally differentiated and stem cells, respectively. Dashed and solid lines correspond to averages after 1 *×* 10^3^ and 7 *×* 10^3^ time steps, respectively. The value of *α* was set at 0.5 (left) and 2 (right). The average was calculated from 5 *×* 10^2^ model realizations. Initial conditions consisted of 20 stem and 20 terminally differentiated cells at the leftmost boundary of the lattice, and all other lattice sites empty. Model parameters were set at *ν* = 0.8, *d* = 0.5, *K*_*S*_ = 32, *K*_*T*_ = 22, 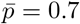. Error bars denote the standard error of the mean.

### 3.3. Feedback strength and local fluctuations control tissue invasive behavior intwo dimensions.s

Next, we computationally studied the dependence of the mean front speed and geometry on the feedback inhibition strength *α*, and on the difference between stem and terminally differentiated cell motility *K*_*s*_ and *K*_*T*_, respectively. We considered a semi-infinite cylindrical lattice, defining a lattice ℒ = ℕ *×* {1, …, *L*,}*L ∈* ℕ, with reflecting boundary conditions at the left boundary and periodic boundary conditions on the top and bottom boundaries, considering no boundary on the right. The initial conditions used consisted of *N* = min {*K*_*S*_, *K*_*T*_} cells within all nodes at the left boundary in both stem and terminally differentiated cell lattices and zero cells in all other nodes in both lattices.

To gauge the front speed, we fixed the simulation time to 3 *×* 10^3^ timesteps and measured the position of the front at the end of the simulation. We defined the front position *F* as the first lattice site up to which at least 95% of the total cell population is contained; mathematically, *F* is a random variable defined as

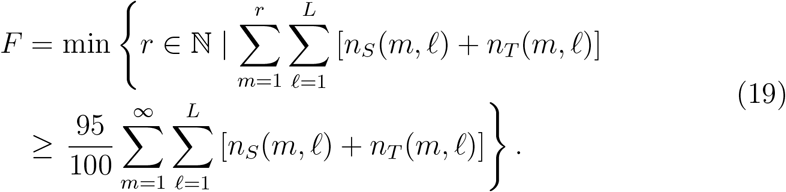

To characterize the irregularity of the invasive front, we identified the first node of each lattice row, where at least 95% of the total row population is found, and joined each of these nodes to the next with a straight line, and measured the length of this envelope. Mathematically, we defined the first front node of each row as

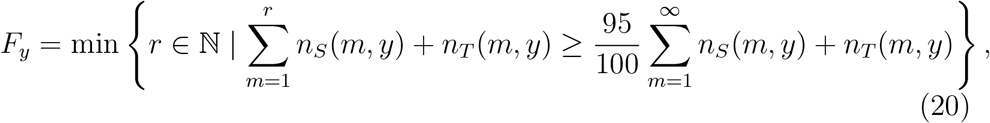

such that the length of the front envelope is given by the random variable

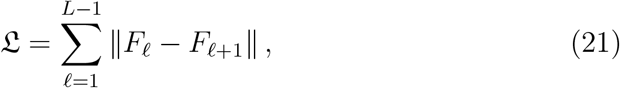

where the distance is the Euclidean distance. Thus, the greater the value of L, the more irregular the front, and vice versa. An example of this measure can be observed in Fig. 7.

**Figure 7.**
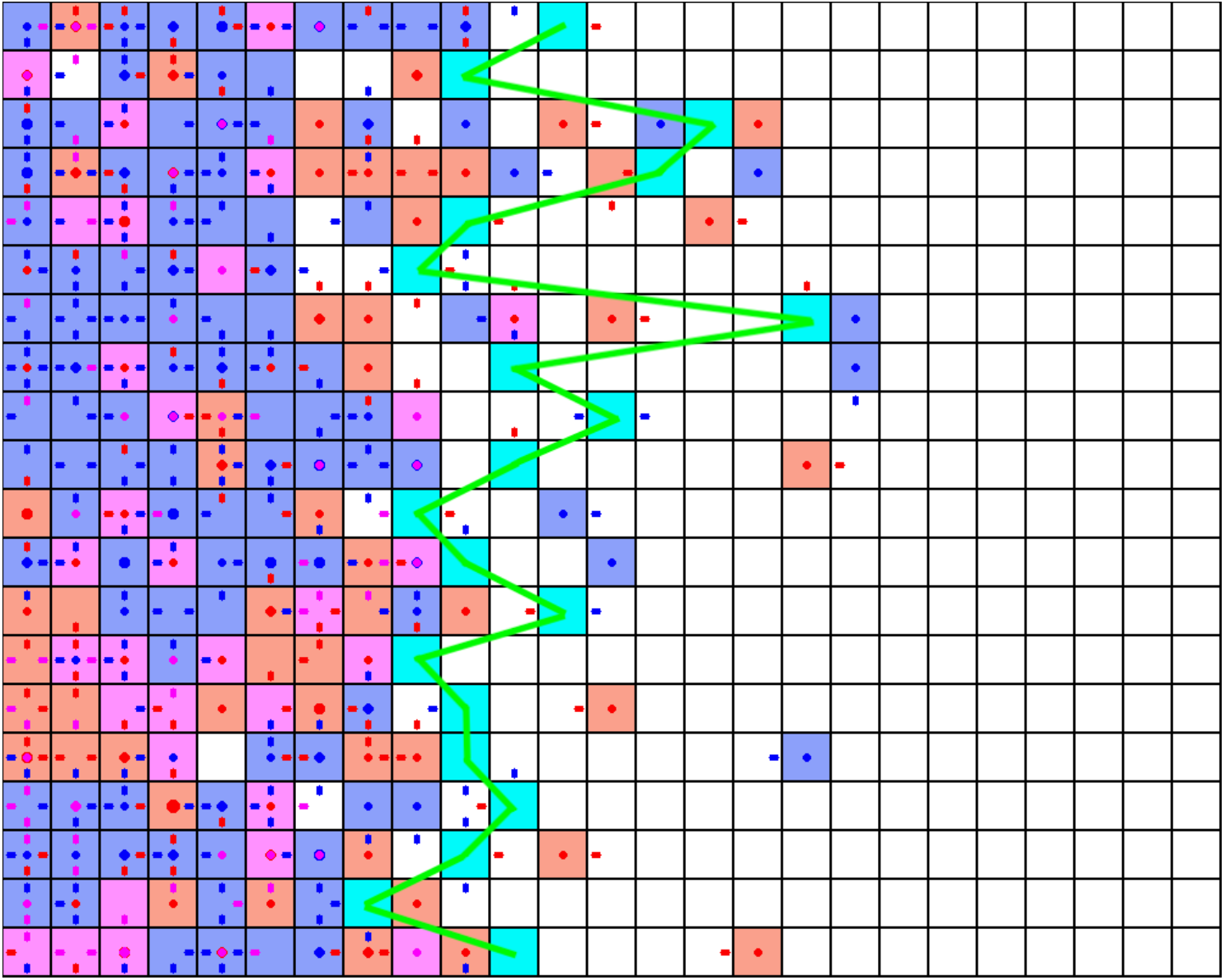
Snapshot of a 2D simulation showing the front envelope. White squares correspond to empty lattice sites. Dark blue squares denote sites with many more CSCs than TDs. Red squares correspond to sites with many more TDs than CSCs. Magenta squares show sites with a great, but equal, number of TDs and CSCs. Dots correspond to immobile (resting) CSCs (blue) or TDs (red), the size of the dot is proportional to the number of resting cells. Arrows correspond to moving CSCs (blue) or TDs (red), pointing to their direction of movement. Cyan squares denote the first lattice site (from left to right) up to where at least 95% of all cells of the row are contained. The green line is the front envelope, connecting all the cyan squares. Its length is represented by L.

As observed in Fig. 8, the front is consistently more regular the less motile stem cells become, since terminally differentiated cells can only die even if they stray too far. However, there appears to be a minimum exactly where *K*_*S*_ = *K*_*T*_ (at *K*_*S*_ = 16 in the figure). This could be explained by the fact that both cell populations will diffuse forward at the same rate when *K*_*S*_ = *K*_*T*_, and therefore will have maximum overlap, such that terminally differentiated cells will efficiently inhibit stem cells. If *K*_*S*_ ≠ *K*_*T*_, then one cell population will be more motile than the other and will probably overtake it, thus lowering the overlap between them and lowering the inhibitory effect.

**Figure 8.**
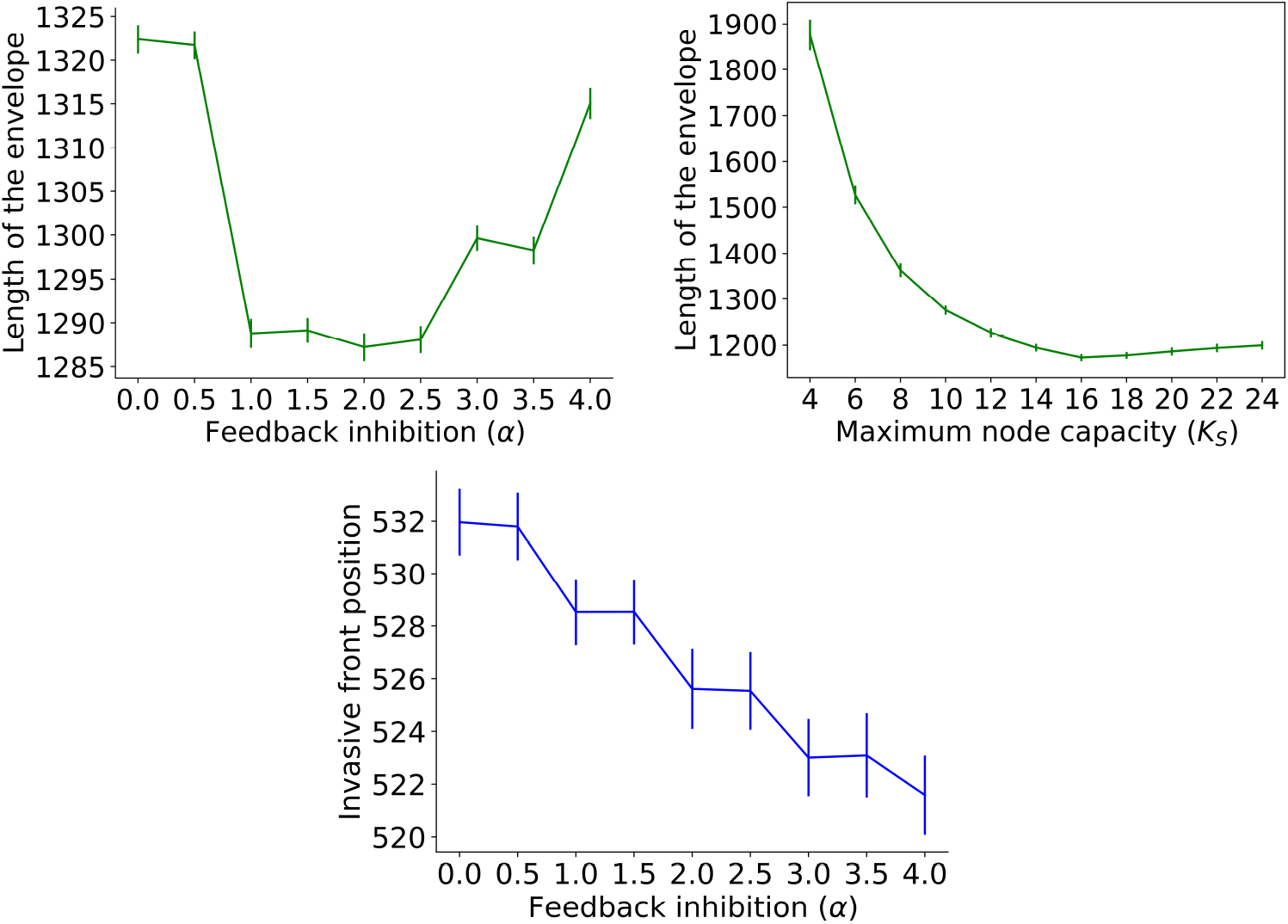
Two-dimensional LGCA simulations. Top: Length of the front envelope L as a function of the feedback inhibition strength *α* (left), and of the maximum number of stem cells per node, *K*_*S*_ (right). Bottom: Position of the front as a function of the feedback inhibition strength *α*. The averages were calculated from 100 model realizations. The values of the observables were recorded after 3 *×* 10^3^ time steps. Unless indicated otherwise, model parameters were set at *L* = 1000, *ν* = 1, *d* = 0.5, *α* = 2, *K*_*S*_ = 8, *K*_*T*_ = 16, 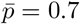. Error bars correspond to one standard deviation.

As opposed to the one-dimensional case, it seems that a higher inhibition strength results in a slower-moving front. This could be explained by an “encapsulation” effect, where the inhibitory effect of terminally differentiated cells decreases the effective division rate of stem cells *ν*, thus decreasing the invasive speed (cp. Eq. 18).

Interestingly, we can investigate the effect of local (node) fluctuations on the system’s invasive behavior. In particular, we can assume node total number of cells fluctuations/variance that 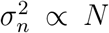, where *n* = *n*_*S*_ + *n*_*T*_, hypothesizing an underlying binomial distribution. From Fig. 8 (top right), we observe that the front envelope decreases as a function of capacity. Since *N* = min {*K*_*S*_, *K*_*T*_ }, a negative monotonic behavior is expected for the front envelope with respect to increasing local fluctuations. Using relation (18), the front speed is inverse proportional to total number of cells and therefore to local noise

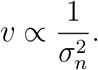

Taking into account these considerations, high local fluctuations lead to unstable fronts that move with lower speed.

Finally, there is a non-monotonic effect of the inhibition strength *α* on the average front regularity L, reaching a minimum at around *α* = 2. Heuristically, when *α* has a low value, stem cells divide independently of the presence of terminally differentiated cells. Thus, if cells can either stay in regions with high cell densities or colonize new sites, they will divide equally, creating a rough invasive front. At intermediate inhibitory levels, stem cells will divide less every time differentiation occurs, or differentiated cells diffuse into their lattice site, keeping growth in check and smoothing out the front. At very high inhibitory levels, however, lattice sites with high numbers of differentiated cells effectively become roadblocks where stem cells cannot divide. Thus, these sites will produce fewer cells and lag behind, while cells with low numbers of differentiated cells will rapidly produce more cells, lengthening the front. This situation would produce a rugged “fingering” pattern, such as that observed in many invasive tumors.

## 4. Discussion

This study highlights the pivotal role of inhibitory feedback in modulating multicellular invasive behavior. Our non-spatial model revealed that differentiated cells’ inhibitory effects significantly reduce stem cell self-renewal, imposing a natural limit on population growth. A bifurcation analysis demonstrated a critical feedback threshold where the population transitions from expansion to extinction, underscoring the importance of inhibitory regulation in maintaining cellular homeostasis. These findings are particularly relevant to wound healing and tissue regeneration, where uncontrolled growth can lead to pathological conditions (34).

Spatial models further elucidated the complexity of these interactions. In the one-dimensional system, the inhibitory feedback was shown not to affect the speed of invasive fronts, but rather the stem/differentiated cell ratio. In two-dimensional systems, the inhibitory feedback impact is more prominent. In particular, the feedback not only dampened the invasion speed but also shaped the morphology of the invasive front. In two-dimensional models, differentiated cells have a greater propensity to become spatially segregated from stem cells, thereby minimizing the extent of inhibitory overlap. This spatial decoupling elucidates the heightened feedback effects on the morphology and velocity of the front observed in two dimensions as opposed to one dimension. High inhibitory feedback resulted in smoother, more regular invasion fronts, whereas low feedback led to irregular, finger-like protrusions reminiscent of aggressive tumor phenotypes (35). The interplay between local motility and inhibitory feedback emerged as a key factor influencing the stability and uniformity of invasive patterns.

Another important aspect is the impact of the node noise/fluctuation, which scales with node capacity. In particular, we observe that high noise reduces tumor speed but increases front instability. The latter is a known phenomenon as noise-driven front instability, as studied by Kessler and Levine (36). Front instabilities, such as fingering, are a characteristic of tissue-invasive behavior. Noise-induced invasions can be biologically relevant during wound healing, spatial spreading of pathological tissues, such as tumors, or bacterial colonies when battling against immune cells.

The effect of inhibitory feedback on front behavior can be associated with a pushed/pulled front transition (37; 38), which impacts their speed and interface stability. In upcoming work, we plan to study in detail whether inhibitory feedback and noise can induce such a transition in lineage-based cell tissue.

Our model’s assumptions limit its scope to simplified stem-differentiated hierarchies. While this simplification may be inadequate for modeling some physiological tissues, where lineages often encompass a long succession of progenitors, in the context of tumors such as glioblastomas (39) or retinoblastomas (40), this process is generally perceived as a single-step transition. Future extensions could easily incorporate multiple lineage states or concurrent cellular events by considering more cell types between the two considered here. Similarly to other models, which consider more intermediate stages (for example, the well-known epidemiological SIR and SEIR models), the inclusion of more intermediate phenotypes would mainly slow down tumor growth and invasive front speed, unless each phenotype is considered to have unique properties and behaviors. Experimental systems such as *in vitro* organoid invasion, epithelial wound healing models, or histological analysis of tumor front morphology could be used to test the predictions of our model regarding feedback and noise-induced front irregularities. Another important limitation/extension of the model is related to the fact that the model omits interactions with immune or stromal cells and environmental heterogeneity. These aspects will be important directions for future work. On the technical side, our model limits updates to one event per site, while in reality cells may act in parallel. On the other hand, this can be easily included in our LGCA by allowing the kinetics operator to be applied more than once. Since our aim was to develop a minimal model that allows analytical tractability, we refrained from this option. Furthermore, since the allowed number of cells per site is limited, applying the kinetics operator would only limit the amplitude of fluctuations in the model that is the model would be less random (41).

These results suggest potential therapeutic strategies for controlling invasive behaviors by targeting differentiation and feedback mechanisms. Enhancing the inhibitory effects of differentiated cells could suppress invasive tendencies in pathological contexts such as cancer. Conversely, modulating feedback strength may optimize regenerative processes in therapeutic applications. Furthermore, increasing the probability of differentiation of cancer stem cells into differentiated cells, while moderately increasing the death rate of differentiated cells, would substantially reduce the proliferative fraction of the tumor, as shown by our simulation results. Future research should aim to incorporate heterogeneity in tissue microenvironments and account for interactions with immune and stromal cells to better reflect biological reality. Additionally, experimental validation of these findings could provide critical insights into the regulatory mechanisms underpinning multicellular invasion.

## Acknowledgments

H.H. would like to acknowledge support from the Volkswagen Stiftung “Life?” initiative (Grant No. 96732), the RIG-2023-051 Khalifa University grant, and the UAE-NIH Collaborative Research grant AJF-NIH-25-KU.

## References

[1] L. Li, Y. He, M. Zhao, and J. Jiang, “Collective cell migration: Implications for wound healing and cancer invasion,” Burns & trauma, vol. 1, no. 1, pp. 2321–3868, 2013.

[2] D. Stichel, A. M. Middleton, B. F. Müller, S. Depner, U. Klingmüller, K. Breuhahn, and F. Matthäus, “An individual-based model for collective cancer cell migration explains speed dynamics and phenotype variability in response to growth factors,” NPJ systems biology and applications, vol. 3, no. 1, p. 5, 2017.

[3] A. Y. Wong and J. L. Whited, “Parallels between wound healing, epimorphic regeneration and solid tumors,” Development, vol. 147, no. 1, p. dev181636, 2020.

[4] M. Deyell, C. S. Garris, and A. M. Laughney, “Cancer metastasis as a non-healing wound,” British journal of cancer, vol. 124, no. 9, pp. 1491– 1502, 2021.

[5] A. Yamamoto, A. E. Doak, and K. J. Cheung, “Orchestration of collective migration and metastasis by tumor cell clusters,” Annual Review of Pathology: Mechanisms of Disease, vol. 18, no. 1, pp. 231–256, 2023.

[6] J. N. Rich, “Cancer stem cells: understanding tumor hierarchy and heterogeneity,” Medicine, vol. 95, no. 1S, pp. S2–S7, 2016.

[7] F. Papaccio, F. Paino, T. Regad, G. Papaccio, V. Desiderio, and V. Tirino, “Concise review: cancer cells, cancer stem cells, and mesenchymal stem cells: influence in cancer development,” Stem cells translational medicine, vol. 6, no. 12, pp. 2115–2125, 2017.

[8] X. Zhang, K. Powell, and L. Li, “Breast cancer stem cells: biomarkers, identification and isolation methods, regulating mechanisms, cellular origin, and beyond,” Cancers, vol. 12, no. 12, p. 3765, 2020.

[9] A. Mohan, R. Raj Rajan, G. Mohan, P. Kollenchery Puthenveettil, and T. T. Maliekal, “Markers and reporters to reveal the hierarchy in heterogeneous cancer stem cells,” Frontiers in cell and developmental biology, vol. 9, p. 668851, 2021.

[10] X. Chu, W. Tian, J. Ning, G. Xiao, Y. Zhou, Z. Wang, Z. Zhai, G. Tanzhu, J. Yang, and R. Zhou, “Cancer stem cells: advances in knowledge and implications for cancer therapy,” Signal Transduction and Targeted Therapy, vol. 9, no. 1, p. 170, 2024.

[11] T. M. Parker, K. Gupta, A. M. Palma, M. Yekelchyk, P. B. Fisher, S. R. Grossman, K. J. Won, E. Madan, E. Moreno, and R. Gogna, “Cell competition in intratumoral and tumor microenvironment interactions,” The EMBO journal, vol. 40, no. 17, p. e107271, 2021.

[12] A. Spinazzola, T. Carvalho, M. A. F. Pinto, M. Marques-Reis, A. Gutiérrez-García, D. Accardi, and E. Moreno, “Heterotypic competition between cancer cells and hepatocytes generates heterogeneous context-dependent phenotypes,” bioRxiv, pp. 2024–05, 2024.

[13] P. Friedl and D. Gilmour, “Collective cell migration in morphogenesis, regeneration and cancer.,” Nature reviews. Molecular cell biology, vol. 10, pp. 445–57, jul 2009.

[14] P. Friedl and R. Mayor, “Tuning collective cell migration by cell-cell junction regulation,” Cold Spring Harbor Perspectives in Biology, vol. 9, no. 4, 2017.

[15] M. J. Bogdan and T. Savin, “Fingering instabilities in tissue invasion: An active fluid model,” Royal Society Open Science, vol. 5, no. 12, 2018.

[16] V. Cristini, H. B. Frieboes, R. Gatenby, S. Caserta, M. Ferrari, and J. Sinek, “Morphologic instability and cancer invasion.,” Clinical cancer research : an official journal of the American Association for Cancer Research, vol. 11, pp. 6772–9, oct 2005.

[17] C. Guiot, P. P. Delsanto, and T. S. Deisboeck, “Morphological instability and cancer invasion: a’splashing water drop’analogy,” Theoretical Biology and Medical Modelling, vol. 4, pp. 1–6, 2007.

[18] C. Jiang, C. Cui, L. Li, and Y. Shao, “The anomalous diffusion of a tumor invading with different surrounding tissues,” PLoS ONE, vol. 9, no. 10, 2014.

[19] A. Sottoriva, L. Vermeulen, and S. Tavaré, “Modeling evolutionary dynamics of epigenetic mutations in hierarchically organized tumors.,” PLoS computational biology, vol. 7, p. e1001132, may 2011.

[20] A. C. Aristotelous and R. Durrett, “Fingering in stochastic growth models,” Experimental mathematics, vol. 23, no. 4, pp. 465–474, 2014.

[21] M. Ben Amar and C. Bianca, “Onset of nonlinearity in a stochastic model for auto-chemotactic advancing epithelia,” Scientific reports, vol. 6, no. 1, p. 33849, 2016.

[22] J. B. McGillen, E. A. Gaffney, N. K. Martin, and P. K. Maini, “A general reaction-diffusion model of acidity in cancer invasion,” Journal of Mathematical Biology, vol. 68, no. 5, pp. 1199–1224, 2014.

[23] E. Gavagnin, M. J. Ford, R. L. Mort, T. Rogers, and C. A. Yates, “The invasion speed of cell migration models with realistic cell cycle time distributions,” Journal of Theoretical Biology, vol. 481, no. June, pp. 91–99, 2019.

[24] N. J. Poplawski, A. Shirinifard, U. Agero, J. S. Gens, M. Swat, and J. a. Glazier, “Front instabilities and invasiveness of simulated 3D avascular tumors.,” PloS one, vol. 5, p. e10641, jan 2010.

[25] H. Youssefpour, X. Li, a. D. Lander, and J. S. Lowengrub, “Multispecies model of cell lineages and feedback control in solid tumors.,” Journal of theoretical biology, vol. 304C, pp. 39–59, mar 2012.

[26] B. Werner, J. G. Scott, A. Sottoriva, A. R. Anderson, A. Traulsen, and P. M. Altrock, “The cancer stem cell fraction in hierarchically organized tumors can be estimated using mathematical modeling and patient-specific treatment trajectories,” Cancer research, vol. 76, no. 7, pp. 1705–1713, 2016.

[27] D. Zhou, Y. Luo, D. Dingli, and A. Traulsen, “The invasion of dedifferentiating cancer cells into hierarchical tissues,” PLoS computational biology, vol. 15, no. 7, p. e1007167, 2019.

[28] P. Uhl, J. Lowengrub, N. Komarova, and D. Wodarz, “Spatial dynamics of feedback and feedforward regulation in cell lineages,” PLoS Computational Biology, vol. 18, no. 5, pp. 1–17, 2022.

[29] K. Böttger, H. Hatzikirou, A. Voss-Böhme, E. A. Cavalcanti-Adam, M. A. Herrero, and A. Deutsch, “An emerging allee effect is critical for tumor initiation and persistence,” PLoS computational biology, vol. 11, no. 9, p. e1004366, 2015.

[30] D. Reher, B. Klink, A. Deutsch, and A. Voss-Böhme, “Cell adhesion heterogeneity reinforces tumour cell dissemination: novel insights from a mathematical model,” Biology direct, vol. 12, pp. 1–17, 2017.

[31] O. Ilina, P. G. Gritsenko, S. Syga, J. Lippoldt, C. A. La Porta, O. Chepizhko, S. Grosser, M. Vullings, G.-J. Bakker, J. Starruß, et al., “Cell–cell adhesion and 3d matrix confinement determine jamming transitions in breast cancer invasion,” Nature cell biology, vol. 22, no. 9, pp. 1103–1115, 2020.

[32] A. Deutsch, J.M. Nava-Sedeño, S. Syga, and H. Hatzikirou, “BIOLGCA: A cellular automaton modelling class for analysing collective cell migration,” PLoS Computational Biology, vol. 17, no. 6, pp. 1–22, 2021.

[33] S. Syga, H. P. Jain, M. Krellner, H. Hatzikirou, and A. Deutsch, “Evolution of phenotypic plasticity leads to tumor heterogeneity with implications for therapy,” PLOS Computational Biology, vol. 20, no. 8, p. e1012003, 2024.

[34] P. Martin and R. Nunan, “Cellular and molecular mechanisms of repair in acute and chronic wound healing,” British Journal of Dermatology, vol. 173, no. 2, pp. 370–378, 2015.

[35] H. B. Frieboes, J. S. Lowengrub, S. Wise, X. Zheng, P. Macklin, E. L. Bearer, and V. Cristini, “Computer simulation of glioma growth and morphology.,” NeuroImage, vol. 37 Suppl 1, pp. S59–70, jan 2007.

[36] D. Kessler and H. Levine, “Fluctuation-induced diffusive instabilities,” Nature, vol. 394, no. 6693, pp. 556–558, 1998.

[37] W. van Saarloos, “Front propagation into unstable states,” Physics Reports, vol. 386, no. 2-6, pp. 29–222, 2003.

[38] J. Garnier, T. Giletti, F. Hamel, and L. Roques, “Inside dynamics of pulled and pushed fronts,” Journal des Mathematiques Pures et Appliquees, vol. 98, no. 4, pp. 428–449, 2012.

[39] M.L. Suvà, E. Rheinbay, S. M. Gillespie, A. P. Patel, H. Wakimoto, S. D. Rabkin, N. Riggi, A. S. Chi, D. P. Cahill, B. V. Nahed, et al., “Reconstructing and reprogramming the tumor-propagating potential of glioblastoma stem-like cells,” Cell, vol. 157, no. 3, pp. 580–594, 2014.

[40] A. Barua, A. Beygi, and H. Hatzikirou, “Close to optimal cell sensing ensures the robustness of tissue differentiation process: the avian photoreceptor mosaic case,” Entropy, vol. 23, no. 7, p. 867, 2021.

[41] S. Syga, J.M. Nava-Sedeño, L. Brusch, and A. Deutsch, “A lattice-gas cellular automaton model for discrete excitable media,” in Spirals and Vortices: In Culture, Nature and Science, pp. 253–264, 2019.

